# Development of Gene-Specific Real-Time PCR Screening Method for Detection of *cp4-epsps* Gene in GM Crops

**DOI:** 10.1101/155127

**Authors:** Solmaz Khosravi, Masoud Tohidfar, Parisa Koobaz

## Abstract

Among the genetically modified (GM) crops that are being approved for commercialization, herbicide resistant crops, especially those harboring *cp4-epsps,* have a considerable contribution. Gene-specific methods can be used to screen the presence of GMOs. To establish an effective qualitative and quantitative screening method, a set of primers were designed considering the *cp4-epsps* sequence. The specificity, the limit of detection, the efficiency, and the ability to quantify the GMO content were tested in GM cotton, soybean, and canola events. The results demonstrated that the primers can specifically detect *cp4-epsps* GM crops. The limit of detection was found to be 0.4 ng /μl DNA per PCR reaction with the ability to detect 1-16 copies of the haploid genome of each GM event. The efficiency of this screening method (which was 94-110 % with an R2 higher than 0.96) indicated that these new primers can be applied to the screening of GM samples that contain the *cp4-epsps* gene. Also, the gene-specific real-time PCR screening method could be successfully developed for qualification of different types of GM cotton, soybean and canola events with the construction of a serial dilution ranging from 10 % to 1 %.

## Introduction

To meet the food demands of the world’s ever-growing population, modern biotechnology has been exploited in agriculture to provide specific crops that can help build the foundation for sustainable food production (*1*). The crops, known as genetically modified organisms (GMOs), were cultured for the first time in 1996 in a cultivation area of 1.7 million hectares. Now, the cultivation area has reached 175.2 million hectares with 27 countries involved in the production (*2*). In 2013, 57 % of this cultivation area belonged to herbicide-tolerant (HT) crops among which glyphosate and glufosinate tolerant varieties were the most common ones. The major HT resistant crops are cotton, canola, soybean, maize, wheat, sugar beet, potato, rice, flax, and alfalfa (*2*). There are several genes such as *gox*, *als*, *bar*, *pat*, and *cp4-epsps* (*3*) that are used to produce HT resistant crops. Among these genes, *cp4-epsps* is the most common gene being used (*4*).

With the production of GM crops and the increasing acceptance of such crops, authorities have made regulations to make it easier to monitor the production and use of these crops on the commercial scale. One such regulation is the labeling of GMOs (*5*).The threshold for unintentional presence of GMOs varies depending on each country’s regulations. For example, it is 0.9 %, 3 %, and 5 % in the EU, Korea and, Japan, respectively (*6*). Australia, New Zeeland, and Brazil have reported 1 % threshold for GM food labeling (*6*). However, for unauthorized GMOs the threshold is zero (*7*).

For this reason, multifarious methods, based on either protein or nucleic acid, have been developed for proper detection of GMOs. The nucleic-acid-based methods include screening, gene, construct, and event-specific methods (*8*). The screening method is usually less reliable due to yielding false positive results (*9*). Since some constructs are used to produce different GMOs, the construct recognition method cannot be specific and accurate enough (*10*). The event-specific screening method uses the unique sequences at the junction of 5′ or 3′ flanking region of genomic DNA and inserted gene. Therefore, the above-mentioned method is very specific and has become the most important method for the identification of GM crops (*11*, *12*). However, the information regarding the junction areas in different GM crops is not always available. Finally, real-time-based systems are widely being used to verify the quantification of GMOs because of their sensitivity, specificity, and also reproducibility of results (*13*). A real-time screening technique was developed to detect *cry1A.105* and *cry2Ab2* genes in genetically modified organisms (*14*). The quantitative PCR screening methods were used for the detection of Roundup Ready, LibertyLink and CryIAb traits in GM products (*15*). Recently, an event-specific detection system for stacked trait maize was introduced (*16*). The detection of soy and maize samples containing *cry1Ab* and *cp4*-*epsps* genes was performed through multiplex PCR (*17*). Also, PCR techniques were employed to detect *35 S*, *Nos* and *cp4*-*epsps* genes in GM soybean and maize (*18*). A multi-target Taq Man real-time PCR was developed for simultaneous detection of different GM crops including cotton, eggplant, maize, potato, rice, and soybean (*19*). More recently, a detection method based on loop-mediated isothermal amplification was used to monitor insect and herbicide resistant crops some of which harbored the *cp4*-*epsps* gene (*20*). The application of this method provided the on-site detection of GM status of samples. However, it is dependent on using three sets of specific primers to detect each gene of interest.

In the present study, we report specific primers designed for the detection of *cp4-epsps* gene in GM crops including soybean, canola, and cotton based on real-time assays. The primers were assayed for their specificity, limit of detection, efficiency, and the ability to quantify the GMO content.

## Materials and Methods

### Sample Material

Certified materials (CRMs) as reference materials harboring *cp4-epsps* gene including 0906-D event MON88913 cotton seed powder, 0906-B event MON89788 soybean powder, and 1011-A event MON88302 Canola were provided by AOCS (American Oil Chemists’ Society, IL, USA). Non-GM materials for each crop were also provided by Seed and Plant Improvement institute, Iran. Also, CRMs lacking the *cp4-epsps* gene, including 0711-A event Ms1 canola and 0306-E2 event LL25 cotton, were prepared to control the specificity of designed primers.

### DNA Extraction

The isolation of DNA from seed powders was carried out according Dellaporta protocol (1983) with some modifications (*21*). Then, all the samples were quantified using a NANO Drop 1000 spectrophotometer (Thermo Scientific, USA).

### Primer Designs

Primers were designed based on the sequence of synthetic construct CP4 EPSPS glyphosate tolerance protein gene available in the National Center for Biotechnology Information (NCBI) with the accession number of JF445290. Using a blastn tool, all the sequences that showed high query coverage with *cp4*-*epsps* (JF445290) were analyzed. The selected sequences from blastn results were multiple aligned using MEGA4 (*22*). Using Vector NTI Software (Advanced 10, Invitrogene), a set of primers were designed from the conserved regions of the *cp4-epsps* gene (Table 1). Moreover, the primers for endogenous genes for each crop according to the public database of GMO detection method database (GMDD) were used to verify the quality of the applied DNA in PCR reactions (Table1).

**Table 1.**
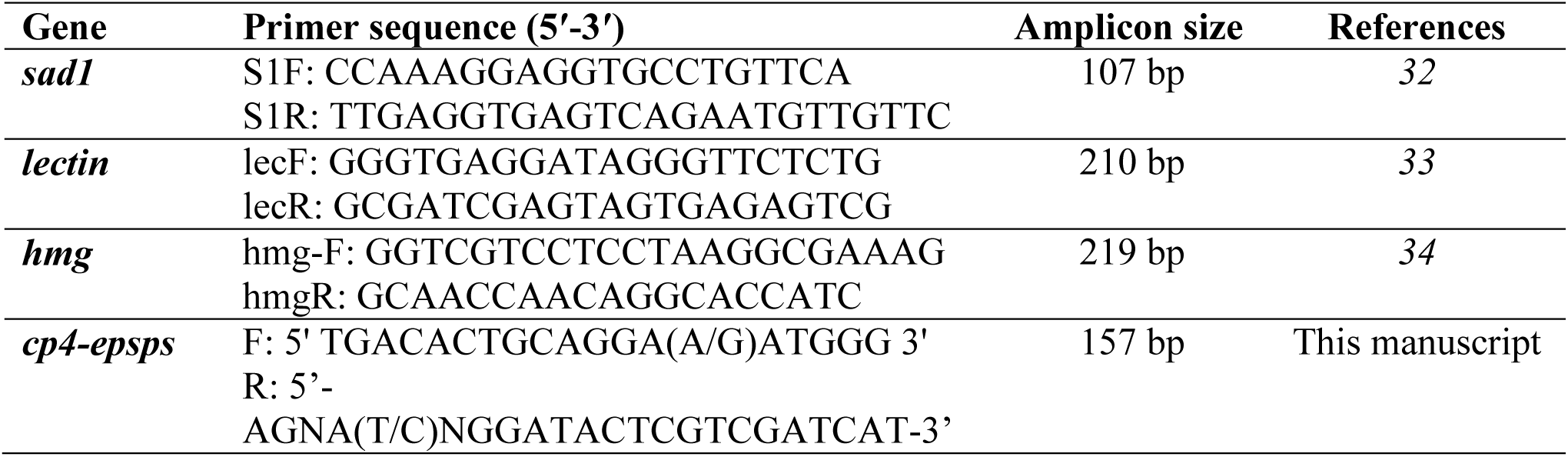
Primer sequences for detection of *cp4-epsps* gene and endogenous genes of cotton (*sad1*), soybean (*lectin*) and canola (*hmg*).

### Polymerase chain reactions

To confirm the presence of the *cp4-epsps* gene, PCR reactions with the total volume of 20 μl containing 2 μl from each of 10 mM dNTPs, 50 mM MgCl2, 10X Taq buffer, 10 pM of forward and reverse primers, 0.3 μl Taq DNA polymerase (5 *u*/μl), and 30 ng of DNA template were performed under the following program: 95 °C 5 min., 35 cycles of 95 °C 30 min., 60 °C 1 min., 72 °C 1 min., and 72 °C 3 min. The PCR reactions to detect the endogenous genes in each crop were also performed under the same conditions. The specificity of the designed primers for the *cp4-epsps* gene was tested by conducting eight PCR runs for CRMs when both containing and lacking the cp4-*epsps* gene.

### Real-time PCR

All real-time PCR reactions for the designed primers to detect *cp4*-*epsps* gene were carried out in 96-well micro-titer plate with the total volume of 25 μl containing 12.5 μl 2X green star qPCR master mix (Bioneer Trade Co., China), 10 pM of forward and reverse primers, 0.5 μl Rox dye, DNA template, and DEPC water. Samples were amplified on MyiQ single color real-time detection system (Bio-Rad, USA) with the following program: 95 °C 5 min., 35 cycles of 95 °C 30 Sec, 60 °C 30 Sec., 72 °C 30 Sec., and 72 °C 3 min.

The efficiency of *cp4*-*epsps* primers was evaluated using real-time PCR analysis by the serial dilution of CRMs MON88913, MON89788, and MON88302. The efficiency test was performed separately for each CRM. The final concentrations used in the analysis were 50, 10, 5, 2, 0.4 and 0.08 *ng*/μl DNA per PCR reaction. The ability of primers to detect the copy number in each CRM was evaluated through dividing the amount of the sample DNA (pg) by the published average 1C value for each GM crop (*23*). To do this, samples containing only GM 1 % with the concentrations of 2 and 0.4 *ng*/μl DNA were applied in real-time assays. These two concentrations were selected based on the lowest concentrations which were detected in the efficiency experiments.

To set a real-time PCR-based quantification system for HT resistant samples harboring *cp4*-*epsps* gene derived from mixed samples, real-time PCR reactions were carried out on different series of CRMs to build a calibration curve based on the logarithm of each CRM content (10, 5, 2, 1 and 0 %) being plotted against ΔCt (Ct_GMO_-Ct_Reference gene_) (*24*). The reference genes including *lectin*, *Sad1,* and *hmg* were used to determine the ΔCt for soybean, cotton, and canola, respectively. The different CRM contents were prepared by mixing the GM and non-GM samples. The equation obtained from this curve was used to determine the unknown percentage of HT resistant samples.

## Results

### Primer Design

Primers were designed based on the sequence of synthetic construct *cp4-epsps* glyphosate tolerance protein gene. By the use of Blastn and MEGA alignment, conserved regions of *cp4-epsps* gene in the available data were selected (Fig. 1). Then, the primers were designed to produce 157 bp amplicons. Finally, the primers were aligned with the partial sequences of GM canola, soybean, and cotton provided by the public database GMDD and also the *cp4-epsps* sequences in the nucleotide sequence collection of NCBI. The query coverage resulting from this alignment was above 90 %.

**Fig. 1.**
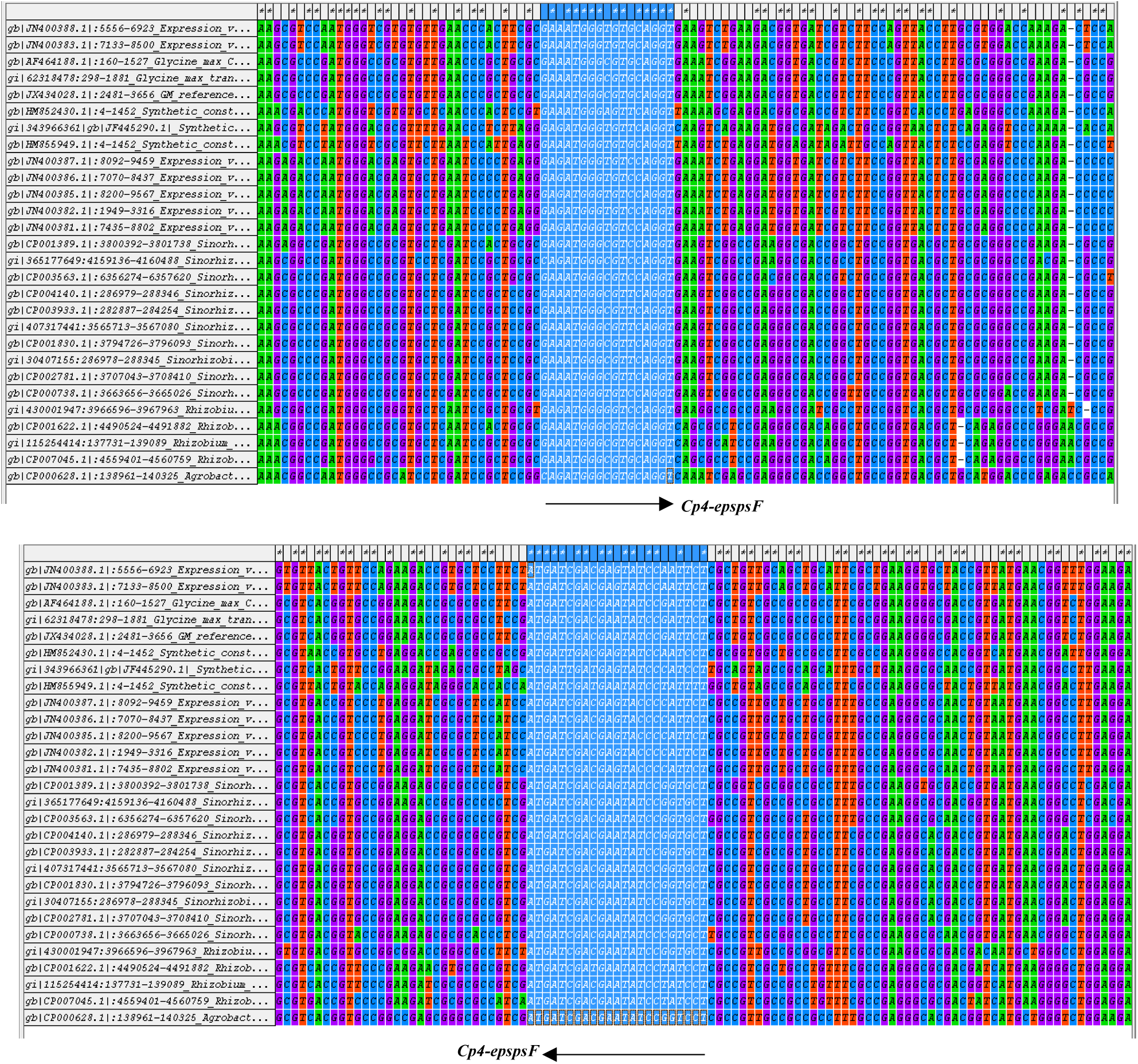
Multiple aligment results on which forward primer (cp4-epspsF) and reverse primer (cp4-epspsR) were desigened for recognition of *cp4*-*epsps* gene in different GM crops.

### Specificity

The endogenous genes including *lectin*, *sad1*, and *hmg* were successfully amplified under the specified condition and could produce amplicon sizes of 210, 107, and 219 bp, respectively (Fig 2). Non-GM cotton, canola, soybean along with MON89788 soybean and Ms1 canola were tested for the *cp4-epsps* gene. The results showed that positive controls MON89788 soybean, MON88913 cotton, and MON88302 canola produced the amplicon size of 157 bp, whereas other tested materials did not show any amplicon (Fig. 3).

**Fig. 2.**
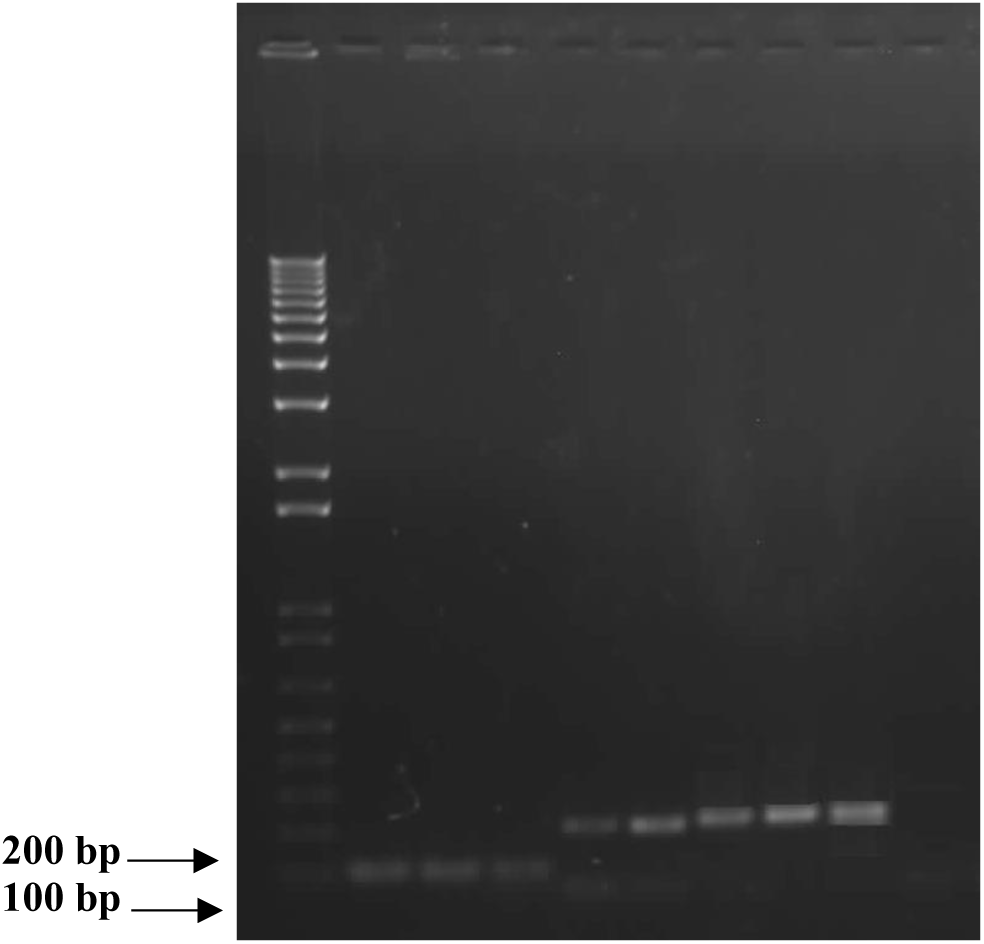
One percent agaros gel electrophoresis of PCR with endogenous primers for each CRM. Lanes 1-3: Ampilication results of *sad1* primers in MON88913, LL25 and non-GM cotton (107 bp). Lanes 4-5: Ampilication results of primers lectin MON89788 and non-GM soybean (210 bp). Lanes 6-8: Ampilication results of *hmg* primers in MON88302, Ms1 and non-GM canola (219 bp). Lane 9: water as negative controls. Lane M: 1Kb Plus DNA ladder

**Fig. 3.**
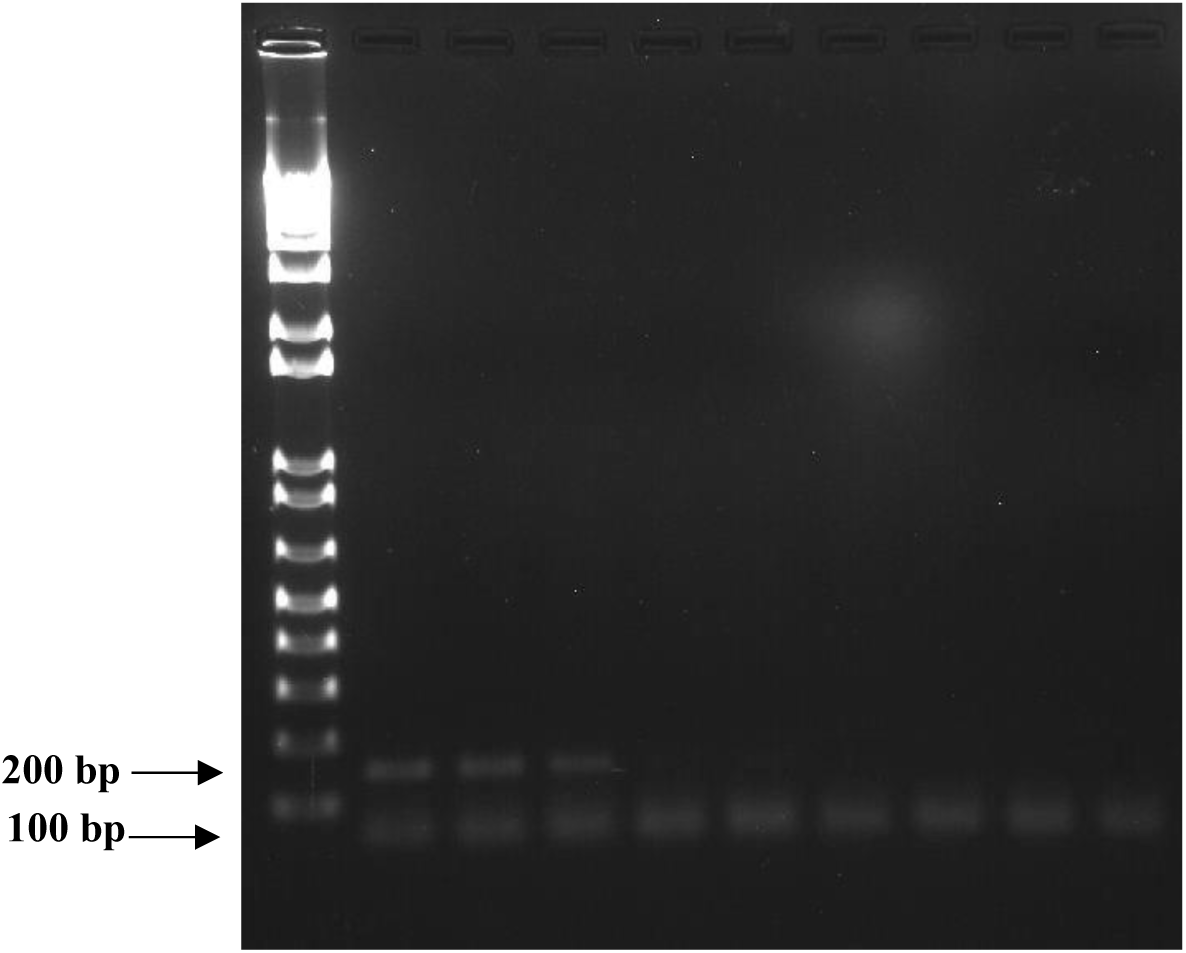
1.5 percent agaros gel electrophoresis of PCR with desigend epspps primers. Lanes 1-3: HT resistant crops MON88913 Cotton, MON89788 Soybean and MON88302 Canola (157 bp), respectively. Lanes 4-5: Ms1and LL25 (GM crops lacking epsps). Lanes 6-8: non-GM cotton, canola and soybean. Lane 9: Water as negative control.. Lane M: 1Kb Plus DNA lad

### Limit of detection and Efficiency

To trace the limit of detection, the different concentrations of 50, 10, 5, 2, 0.4 and 0.08 *ng* /μl in each real-time PCR reaction were used for each reference material. The lowest concentration that was detected in each sample was 0.4 (*ng*/μl) (Table 2). The Ct values for MON88302, MON88913, and MON89788 were 32.26, 30.81, and 32.27, respectively.

**Table 2.** Limit of detection for *cp4*-*epsps* gene in MON89788 soybean, MON88913 cottonseed and MON88302 Canola.

| Amount of<br>DNA ( ng /μl ) | MON89788 Soybean | MON88913<br>Cottonseed | MON88302 Canola |
| --- | --- | --- | --- |
|  | Ct mean |  |  |
| 50 | 25.11± 0.08 | 23.24± 0.3 | 25.38± 0.12 |
| 10 | 27.01± 0.3 | 25.41± 0.21 | 28.45± 0.18 |
| 2 | 28.47± 0.44 | 27.12± 0.1 | 31.95± 0.08 |
| 0.4 | 32.27± 0.43 | 30.81± 0.39 | 32.26± 0.28 |
| 0.08 | NA | NA | N/A |

The efficiency of primers was evaluated for all three CRMs. According to the results, a linear trend was observed for all experiments when Ct values were plotted against the logarithm value of the DNA amount. However, 50 *ng*/μl DNA samples showed out of range results (Fig.4a, b and c). Then, they were discarded from standard curves calculations. The efficiency values for different CRMs varied from 94 to 101 %, and R2 was higher than 0.96.

**Fig 4.**
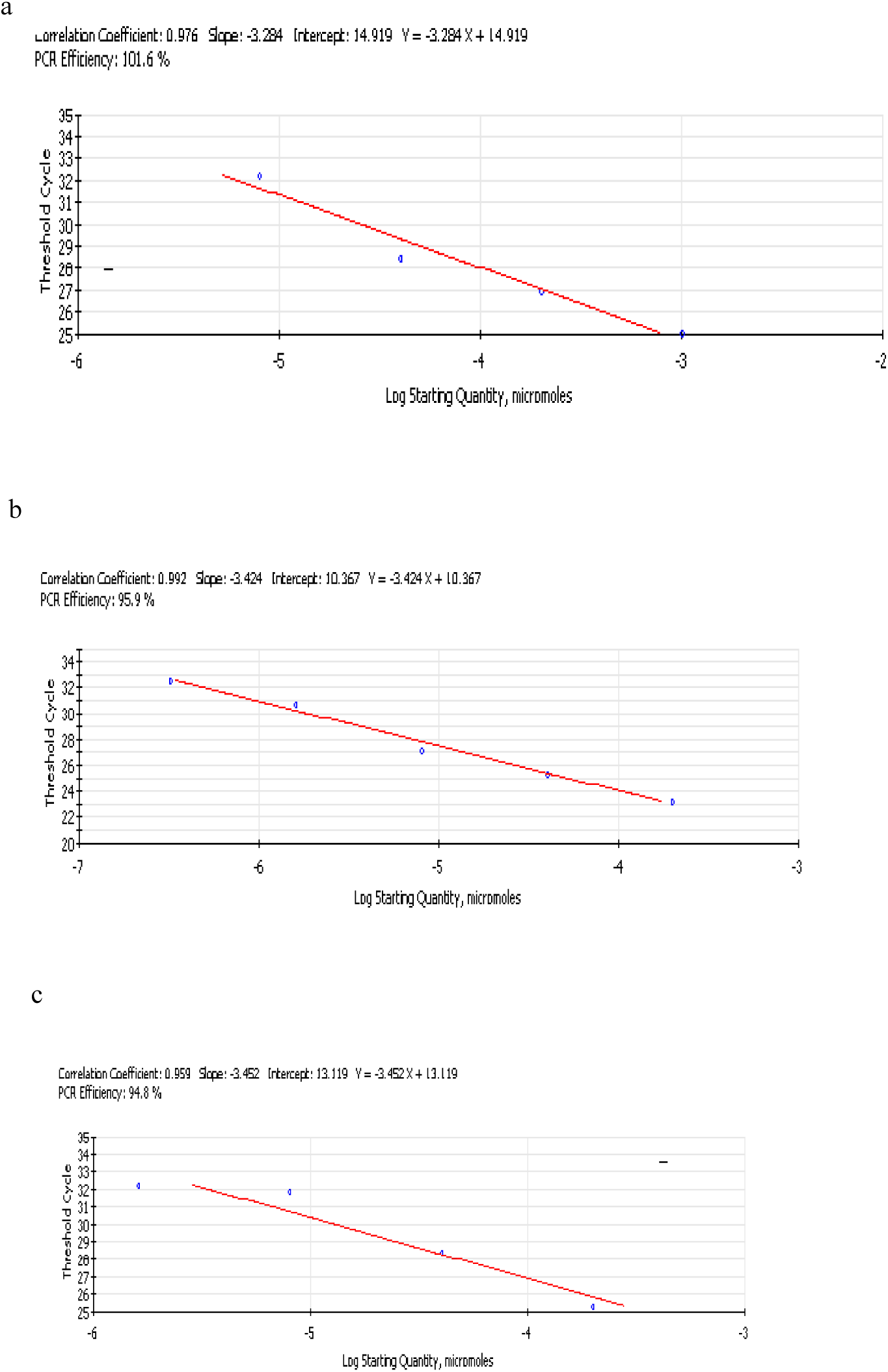
Linear regression of *cp4*-*epsps* gene resulted from serial dilution of 10, 5, 2, 0.4 and 0.08 ng /μl DNA per real-time PCR reaction for a. MON89788 Soybean, b. MON88913 Cottonseed and c. MON88302 Canola

Ct mean values from real-time PCR assays in which the 1 % GM materials in concentrations of 2 and 0.4 *ng*/μl were used are presented for MON88302, MON88913, and MON89788 in Table 3. The copy number of *cp4-epsps* gene in each GM crop was calculated by dividing the DNA amount (pg) by the published average 1C value of 2.33 pg, 1.15 pg, and 1.25 pg for cotton, canola and soybean, respectively (*25*). The copy number calculated by using the sample DNA of 2 and 0.4 *ng*/μl from 1 % GMO was 8 to 1 copies of the haploid genome of GM cotton MON88913. For canola, it was 4 to 16 copies of the haploid genome of GM canola MON88302, and it was 3 to 16 copies of the haploid genome of GM soybean MON89788.

**Table 3.** Estimated copy numbers of *cp4*-*epsps* gene in different 1 % CRMs.

| Event | Amount of<br>DNA ( ng /μl ) | Ct mean | Estimated Copy number per<br>PCR reaction |
| --- | --- | --- | --- |
| MON89788 Soybean | 2 | 30.24± 0.1 | 16 |
| | 0.4 | 32.16± 0.15 | 3 |
| MON88913<br>Cottonseed | 2 | 27.21± 0.13 | 8 |
| | 0.4 | 29.84± 0.08 | 1 |
| MON88302 Canola | 2 | 28.63± 0.09 | 16 |
| | 0.4 | 30.94± 0.15 | 4 |

### GMO quantification

Following the optimization of PCR reactions to detect the *cp4*-*epsps* gene in different GM crops, real-time PCR reactions were completed to quantify the GM content in various samples. Reliability of experiments was confirmed by repeating the reactions for three times. The average Ct values of *cp4*-*epsps* gene are presented in Table 4. The Ct values correlated with the amount of GM content used in each assay. By increasing the GM content from 1 % to 10 %, the Ct values followed a decreasing pattern. This confirms the presence of higher GM content in each mixed sample. Moreover, the samples lacking any GM (0 %) did not produce Ct values.

**Table 4.** The Ct means values of MON88913, MON88302 and MON89788 from real-time assays for *cp4*-*epsps* and reference genes relating each crop.

| GM content (%) | Ct mean value |  |  |
| --- | --- | --- | --- |
|  | MON88913 | MON88302 | MON89788 |
| *cp4-epsps* |  |  |  |
| 0 | - | - | - |
| 1 | 29.65± 0.12 | 33.13± 0.11 | 33.18± 0.15 |
| 2 | 29.28± 0.15 | 32.19± 0.15 | 32.31± 0.13 |
| 5 | 27.09± 0.08 | 30.88± 0.13 | 31.99± 0.08 |
| 10 | 26.37± 0.10 | 29.63± 0.07 | 31.09± 0.05 |
| Reference genes |  |  |  |
|  | *Sad1* | *hmg* | *lectin* |
| 0 | 25.41± 0.45 | 24.71± 0.28 | 22.68± 0.08 |
| 1 | 25.27± 0.3 | 24.73± 0.24 | 22.70± 0.13 |
| 2 | 25.46± 0.27 | 24.73± 0.1 | 22.53± 0.2 |
| 5 | 25.42± 0.03 | 24.72± 0.2 | 22.89± 0.33 |
| 10 | 25.34± 0.39 | 24.73± 0.3 | 22.53± 0.15 |

Once the logarithm of each concentration was plotted against ΔCt, an equation was established upon which the GM content would become estimable (Fig 5a, b and c). The equations resulting from these calibration curves showed an R2 of 0.96-0.99.

**Fig. 5.**
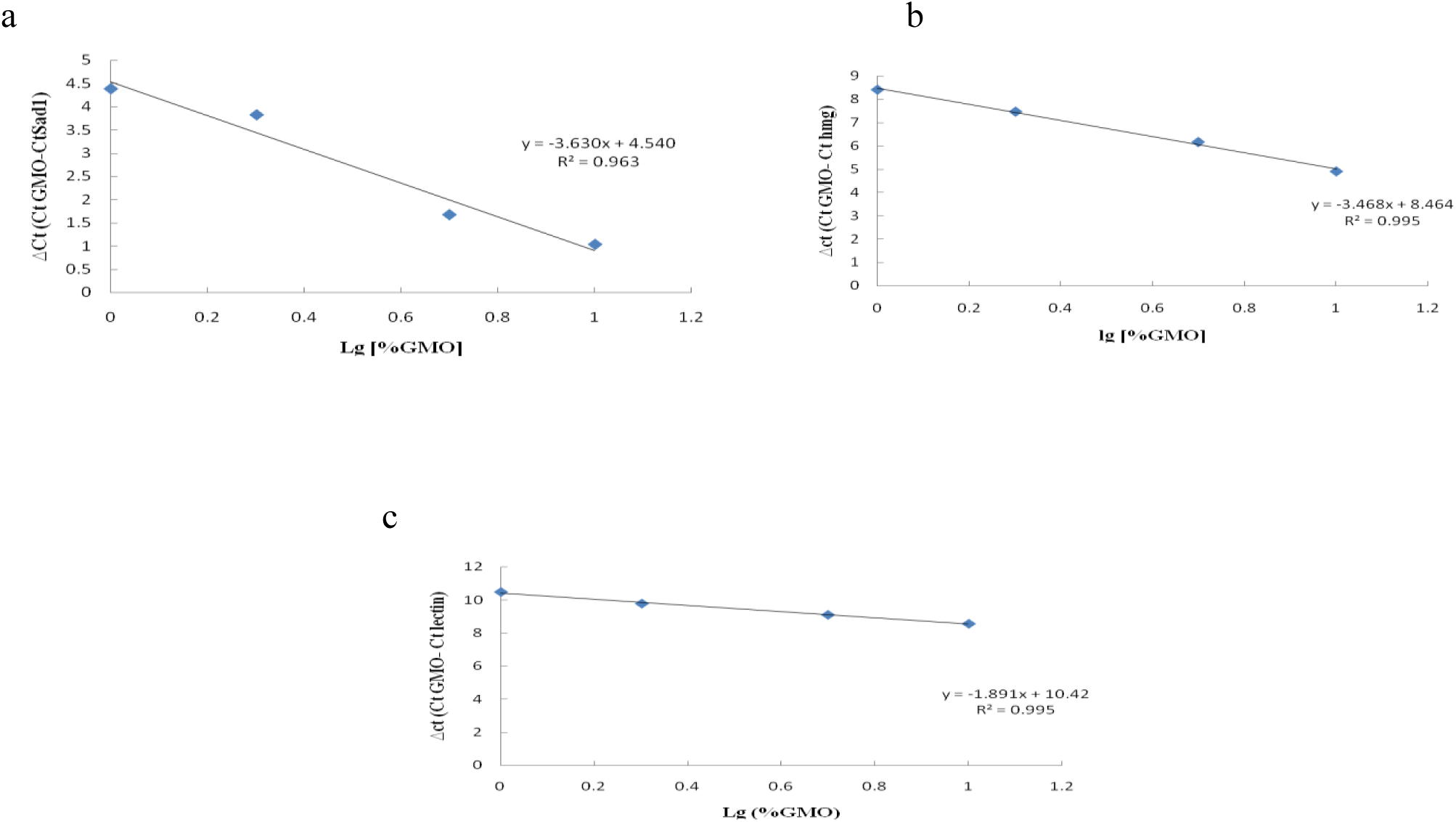
Quantification equations of GM content using real-time PCR for a. MON88913 Cottonseed, b. MON88302 Canola and c. MON89788 Soybean.

## Discussion

The specificity of primers that are used in GM detection programs is of great importance. Based on our bioinformatics analysis, conserved regions of *cp4*-*epsps* sequences in cotton, soybean, and canola and other DNA relevant data were retrieved. The designed primer pair based on conserved regions could discriminate between non-GM crops and GM crops harboring *cp4*-*epsps* with no cross-reactivity, though GM crops containing *bar* were not detected. Furthermore, a set of primers that could specifically identify the GM maize events harboring *cry1A.105* and *cry2ab2* was developed (*14*). PCR primer sets that could amplify *cry2Ab* (326 bp), *P35s* (180 bp), *T-Nos* (195 bp), and *Npt-II* (215 bp) in different crops were also developed (*26*).

The lowest amount that can be reliably detected in a sample is referred to as limit of detection (LOD). The LOD between 0.05 and 0.01 (*ng*/μl) DNA was detected for *cry1A.105* and *cry2ab2* when a serial dilution of 100 % CRM MON 89034 was targeted in real-time assays, which corresponds to 18 to 4 copies (*14*). Recently, different DNA-based detection technologies have been developed to evaluate the copy number of *cp4*-*epsps* gene in different crops (*20*, *19*, *15*). In a multi-target system which was developed for the simultaneous detection of 47 targets, the system was capable of detecting 42, 6, and 88 copies of the *cp4-epsps* gene in cotton MON 88913, cotton MON 71800, and soybean GTS 40-3-2, respectively where the sensitivity of detection was lower than the method used in the present research (*19*). In another study, the lowest copy number which was detectable based on loop-mediated isothermal amplification method was 4 copies of *cp4*-*epsps* (*20*). In addition, the LOD of 2 copy numbers for *cp4-epsps* in soybean GTS 40-3-2 was reported. The established method for detection was based on coSYPS (*15*). The detected LOD in our experiments was 0.4 ng/μl DNA per PCR reaction when the 100 % of the DNA input was GM. The LOD of 0.4 ng /μl DNA was also confirmed when the applied DNA contained just 1 % GM content, which corresponds to 1, 3, and 4 copies of the haploid genome of GM cotton, canola, and soybean, respectively. Therefore, the findings of our research are in accordance with those of Barbue and Singh. The defined threshold for the labeling of GM products in EU is 0.9 % (*27*) which is the strictest among other countries.

The designed *cp4*-epsps primers produced a linear quantitative curve in this research. However, the 50 ng /μl DNA per PCR reaction gave an out of range result that disturbed the efficiency of this method, so it was omitted from efficiency calculations. The PCR efficiency of all experiments with a serial dilution of 10, 5, 2, 0.4, and 0.08 (*ng*/μl) DNA in real time PCR reaction was between 94 % and 101 %, which is in accordance with the European Network of GMO Laboratories (ENGL) criteria (LOD below 20 target copies with PCR efficiency between 89.6 % and 110.2 %) (*27*).

Not only the detection of specific inserted genes into crops but also the quantification of them is a concern today among authorities who supervise the production and distribution of GM crops. In our study, a standard curve for the mixture of non-GM and HT samples containing *cp4*-*epsps* was established. The presence of square regression coefficients (R2) which were higher than 0.96 for *cp4*-*epsps* gene showed this method is suitable for quantitative measurements. Calculation of GM content percentage based on the presented procedure was performed for MON863 which contains *Cry3Bb1* (*28*). However, this method was based on screening and construct specific method, and the detectable LOD was 8 copies in haploid genome.

Also, the quantification of GM tomato Huafan No. 1 which contains anti-sense ethylene-forming enzyme (*EFE*) gene developed based on event specific primers (*29*). According to our experiments, the estimated GM content based on the establishment of standard curve might show a slight variation with the real content. This variation was also observed in other reports (*28*; *29*; *30*). Peccoud and Jacob (1996) reported that molecular fluctuations during real-time assays are associated with applied low copy number of template and varying amplification efficiencies. Therefore, the uncertainty associated with real-time quantification methods may occur due to these factors. Actually, when low levels of DNA from GM samples are targeted to quantitative detection methods, such variations in the measured results are probable and acceptable (*31*).

## Conclusion

We developed an efficient and applicable real-time PCR method for the identification of various GM products including canola, soybean, and cotton based on designing new specific *cp4-epsps* primers. The primers were able to detect both types of *cp4-epsps* genes (type I and II) that exist in different crops.

## Acknowledgment

The authors are grateful to Agriculture Biotechnology Research Institute of Iran (ABRII) for research facilities.

